# TranSynergy: Mechanism-Driven Interpretable Deep Neural Network for the Synergistic Prediction and Pathway Deconvolution of Drug Combinations

**DOI:** 10.1101/2020.07.08.193904

**Authors:** Qiao Liu, Lei Xie

## Abstract

**Motivation:** Drug combinations have demonstrated great potential in cancer treatments. They alleviate drug resistance and improve therapeutic efficacy. With the fast-growing number of anti-cancer drugs, the experimental investigation of all drug combinations is costly and time-consuming. Computational techniques can improve the efficiency of drug combination screening. Despite recent advances in applying machine learning to synergistic drug combinations prediction, several challenges remain. First, the performance of existing methods is suboptimal. There is still much space for improvement. Second, biological knowledge has not been fully incorporated into the model. Finally, many models are lack of interpretability, limiting their clinical applications.

**Results:** We develop a knowledge-enabled and self-attention boosted deep learning model, TranSynergy, to improve the performance and interpretability of synergistic drug combinations prediction. TranSynergy is well designed such that cellular effect of drug actions can be explicitly modeled through cell-line gene dependency, gene-gene interaction, and genome-wide drug-target interaction. A novel Shapley Additive Gene Set Enrichment Analysis (SA-GSEA) method is developed to deconvolute biological pathways that contribute to the synergistic drug combination and improve model interpretability. Extensive benchmark studies demonstrate that TranSynergy significantly outperforms the state-of-the-art method, suggesting the potential of mechanism-driven machine learning. Novel pathways that are associated with the synergistic combinations are revealed and supported by experimental evidence. They may provide new insights into identifying biomarkers for precision medicine and discovering new anti-cancer therapies. Several new synergistic drug combinations are predicted with high confidence for ovarian cancer which has few treatment options.

**Availability:** The code is available at https://github.com/qiaoliuhub/drug_combination

## 1 Introduction

With an increasing comprehensive understanding of the disorder in cancer cells, many anti-cancer drugs are under investigation. However, drug monotherapy suffers limited efficiency due to inherent or acquired resistance (Housman, et al., 2014; Mansoori, et al., 2017; Nikolaou, et al., 2018). Drug combination therapy is an effective strategy to solve this challenging problem (Bayat Mokhtari, et al., 2017; Fitzgerald, et al., 2006; Lehar, et al., 2009; Raghavendra, et al., 2018; Ramsay, et al., 2018). In addition to cancer, synergistic drug combinations have several successful applications in the treatment for other diseases, such as AIDS (Henkel, 1999; Murphy, et al., 2008), and fungal or bacterial infections (Chen, et al., 2016; Groll and Tragiannidis, 2009; Tamma, et al., 2012). Thus, the selection of efficient drug combination therapy for pathogens emerges as a compelling treatment strategy. Considering that the number of anti-cancer drugs has increased drastically, the possible combinations of all these drugs also become enormous (Ali, et al., 2012; Falzone, et al., 2018). Existing experimental method requires a large number of samples with different drug doses and cancer cells (Mott, et al., 2015), thus is infeasible to exhaust all the possible drug combinations. The computational method could be used to pre-select drug combinations with high synergy more cheaply and efficiently. The recent advancement of computational modeling, especially the deep learning technique, has dramatically increased the prediction power of computational models and has many promising applications in the biomedical field. The combination of computational and experimental methods can improve the effectiveness of the drug combination discovery.

The deep learning model has shown better performance than many conventional machine learning algorithms in many biomedical applications (Kalinin, et al., 2018; Mater and Coote, 2019). High-quality experimental drug combination datasets are necessary for the success of deep learning. With the advancement of high throughput drug combination screening tests, the number of samples grows fast so that the data size limitation is considerably alleviated (Bulusu, et al., 2016; Holbeck, et al., 2017; Li, et al., 2018; Menden, et al., 2019; O’Neil, et al., 2016). DeepSynergy is a state-of-the-art deep learning-based prediction model for the prediction of synergistic drug combination. It was trained using the dataset released by Merck (Preuer, et al., 2018). In addition to suboptimal performance, the issue for this model is that the interpretation of model is limited by the way adopted to represent drugs and cell lines as well as the model architecture. For instance, it is difficult to associate the contributions or feature importance of the drug descriptors, including toxophores, physicochemical properties, and fingerprints, with the mechanism of drug action in cells using a feedforward neural network (Benitez, et al., 1997; Castelvecchi, 2016; Challen, et al., 2019).

Recent studies have shown that gene-gene interacting network properties play a critical role in the synergistic drug combination (Cheng, et al., 2019). In addition, cell line drug sensitivity strongly depends on whether the drug directly or indirectly inhibits the essential gene of the cell line (Menden, et al., 2019). Thus, it is desirable to incorporate information from gene-gene interacting network, gene dependency, and drug-target interaction into the deep learning model. To this end, we implemented a mechanism-driven and self-attention boosted deep learning model TranSynergy for the prediction of synergistic drug combinations and the deconvolution of cellular mechanisms contributing to them. Instead of using chemical information as the representation for drugs, we applied the random walk with restart algorithm on a protein-protein interaction (PPI) network to infer a novel drug-target profile as the drug features. For each cell line, we used gene expression, gene dependencies, or PPI derived NetExpress scores that are correlated with the gene dependency as cell line descriptors (Jiang, et al., 2015; Liu, et al., 2019). These mechanism related features make the model readily interpretable. Furthermore, we applied the self-attention to encode the gene-gene interactions responsible for the synergistic drug combination. Attention mechanisms have been widely used in image processing and natural language processing (Bahdanau, et al., 2014; Luong, et al., 2015; Vaswani, et al., 2017) as well as shown promise in the predive modeling of nucleic acid sequences (Liu, et al., 2019). When combining the self-attention encoded representation of drug-target interaction, gene dependency, and NetExpress score, TranSynergy significantly outperforms the state-of-the-art model. To reveal novel pathways that are associated with the synergistic drug combination from the learned biological relations in TranSynergy model, we developed a novel Shapley Additive Gene Set Enrichment Analysis (SA-GSEA) based on SHAP (Lundberg and Lee, 2017). The revealed novel pathway may serve as a patient-specific biomarker for precision medicine or drug targets for discovering new cancer combination therapy. We further applied the model for the prediction of novel synergistic drug combination targeting cancers that have few treatment options. Given the emergence of the next-generation sequencing technology, transcriptome of patient-derived cancer cells can be readily obtained (Buermans and den Dunnen, 2014). The TranSynergy can be used to predict and interpret the synergistic drug combination in distinct patient-derived cancer cells. Our study shows the potential of mechanism-driven interpretable machine learning model in the application of personalized cancer treatment.

## 2 Methods

### 2.1 Drug combination synergy score dataset

The large-scale drug combination screening dataset was initially published by Merck (O’Neil, et al., 2016) and preprocessed to calculate the synergy scores (Preuer, et al., 2018). The screening test was performed with 38 drugs and 39 cancer cell lines. In total, 583 pairs of drug combinations were investigated and 23062 data points were collected. Among them, 2 drugs don’t have any DNA targets and the gene dependencies and gene expression data of four cells are not available. These drugs and cell lines were excluded from our data set. We finally selected 36 drugs that targeted at least one protein and 35 cell lines. The other. The final dataset has 18623 data points and 525 pairs of drug combinations.

### 2.2 Drug representation

Observed drug target profile was collected from two datasets, Drugbank and ChEMBL for the 36 drugs (Gaulton, et al., 2017; Wishart, et al., 2018). The observed drug target matrix is a 36*2401 binary matrix that indicates whether a drug targets a protein. Observed drug target profile was processed with random walk with restart algorithm to obtain a novel drug target profile. We solving the random walk with restart problem with the Fast RWR methods (Tong, et al., 2006). Following is the formal equation:

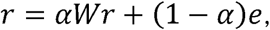

*α* is a hyperparameter equal to 1 - restart rate. We used the restart rate of 0.5. W is the transition matrix denoting the transition probability between nodes. We used the protein-protein interaction network matrix from STRING as the transition probability matrix (Szklarczyk, et al., 2019). The edge weight between nodes is the protein-protein association confidence score. *e* is the seed vector in this equation. We used a drug target binary vector for each drug. 1 denotes that the drug is targeting the protein. *r* is the final probability distribution of each node in the network. Intuitively, *r*[i] denotes the effect of each drug on each protein.

### 2.3 cell line representation

Gene expression profiles were downloaded from Harmonizome (Rouillard, et al., 2016) and were initially collected in Broad Institute Cancer Cell Line Encyclopedia (CCLE) and Genomics of Drug Sensitivity in Cancer (GDSC) (Barretina, et al., 2012; Iorio, et al., 2016; Yang, et al., 2013). Gene dependencies profile dataset is a combined dataset from the Broad Institute Project Achilles (Broad, 2020; Dempster, et al., 2019; Meyers, et al., 2017) and Sanger CRISPR data from Wellcome Trust Sanger Institute, Broad Institute(Behan, et al., 2019; Broad, 2019). The dataset was downloaded from the DepMap portal (McFarland, et al., 2018). Linear regression imputation method was used to fill in the missing value in the dataset with MICE package (van Buuren and Groothuis-Oudshoorn, 2011). NetExpress scores were calculated with the Network analysis of gene essentiality (NEST) tool (Jiang, et al., 2015). The input gene expression was collected as mentioned above and the protein-protein interaction network is downloaded from the STRING database (Szklarczyk, et al., 2019).

### 2.4 Model evaluation

The model evaluation method used is leave-drug-combination-out. The data was split into five folds. The drug pair in one fold does not overlap with the drug pairs in other folds. We used the same training and test split and the same five-folds as those in deepSynergy. Because the drug combination of drug A with drug B and drug B with drug A should have the same drug combination synergy score, the size of training data was doubled by swapping the drug A and drug B. We used mean squared error as the training loss, Spearman correlation and Pearson correlation as evaluation metrics.

### 2.5 Shapley Additive Gene Set Enrichment Analysis

We use the GradientExplainer and DeepExplainer in the SHAP package to calculate Shapley value that characterizes the contribution of each input feature to the prediction model (Lundberg and Lee, 2017). We use k-means to summarize the total dataset as the background dataset. The final Shapley value of each input feature was the average value of 10 tests for each sample data. We then ranked genes based on the Shapley values for the gene-wise features of cell line representation and conducted gene set enrichment analysis to unveil the enriched gene sets with GSEA (Liberzon, et al., 2011; Subramanian, et al., 2005).

## 3 Results

### 3.1 Integration of protein-protein interaction network with drug target information and gene expression information for the representation of drugs and cell lines

The input of our deep learning model includes the vector representations of two drug molecules in the drug combination and cell line that is treated by the drug combination. One popular strategy is to use the physicochemical properties, fingerprints, or toxicophores that are derived from the molecular structure of chemical compound for the representation of drug(Preuer, et al., 2018). There are two disadvantages in using chemical structure as the feature. First, it is not straightforward to establish causal relationship between the physiochemical properties of drugs and the cellular mechanism of drug action. Second, the model is less generalizable when applied to compounds that are not similar to those in the training set (Ayed, et al., 2019). Biological representation based on drug-target interaction profile is an alternative strategy to infer the drug representation vector (Ayed, et al., 2019). Drug targets information that are collected from databases, including Drugbank and ChEMBL (Gaulton, et al., 2017; Wishart, et al., 2018), are mainly proteins that are directly interacted with drugs. We also need to encode the effect of drugs on down-stream non-target proteins and whole biological system. We utilize protein-protein interaction network to infer the drug response of the non-target proteins, since the protein-protein interaction mediates information transmission in the biological system. We apply the random walk with restart algorithm to simulate this network propagation process (Figure 1). Compared with the chemical-based approaches for drug representation, the target-based representation of drug molecules has several advantages. Firstly, drug target information is closely related to the cellular response to the drug treatment at both molecular level and system level. Secondly, it makes it possible to explain the computational model output, drug combination synergy, in terms of the contribution of certain proteins or genes to cancer cells.

**Figure 1.**
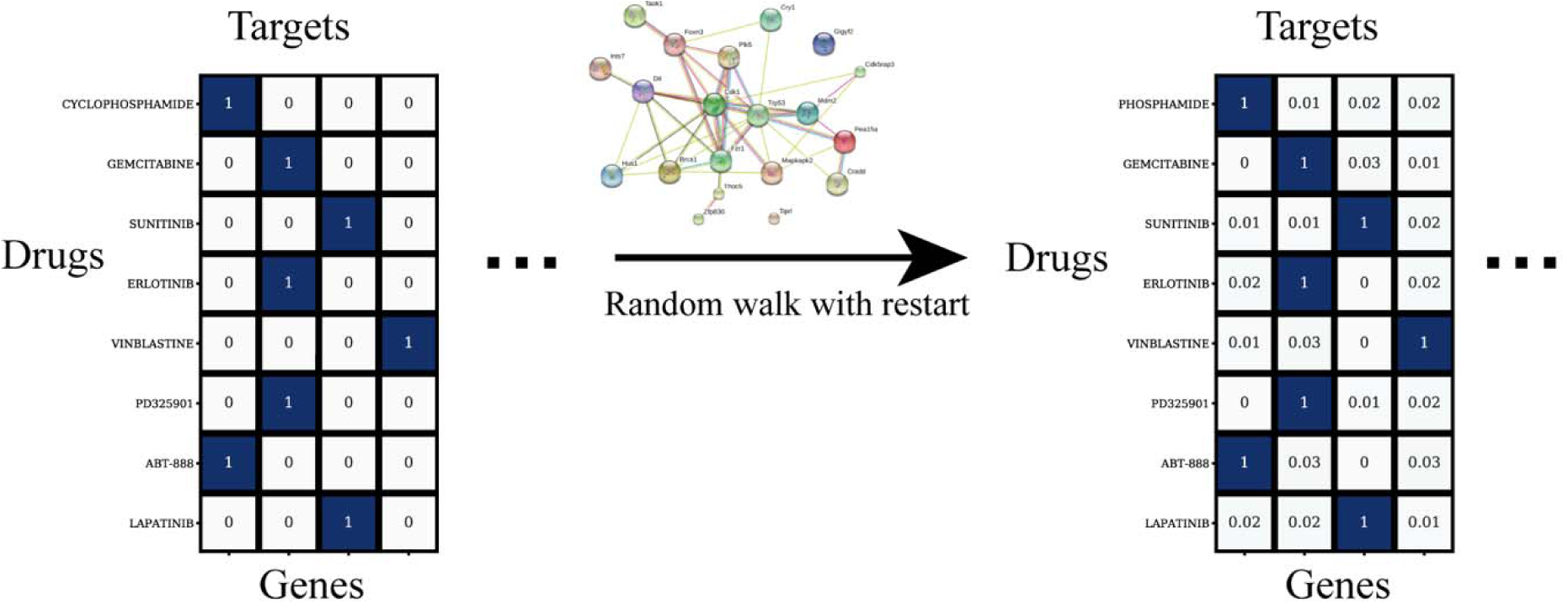
Illustration of genome-wide drug-target profile. Observed drug target profile is processed with random walk with restart to infer drug effects on both targets and non-target proteins.

Another component of inputs is the cell line vector representation because drug-combination has a cell line-specific response. The deepSynergy uses gene expression profile only (Preuer, et al., 2018). We applied a novel alternative strategy to infer cell line vector representation. The essentiality of genes varies in the different cell lines and plays a critical role in the anti-cancer drug sensitivity. Intuitively, drugs that affect more essential proteins will cause the cell to have a more devastating response. Two approaches are adopted to determine gene dependence information. The first one is collected by the BROAD institution using an experimental method (McFarland, et al., 2018). They performed a genome-wide loss-of-function screening with pooled RNAi or CRISPR library and then investigated the resulted cellular response. The second method is through a computational tool that integrates gene expression and gene-gene interaction network information (Jiang, et al., 2015). The calculated scores, called NetExpress score, are shown to indicate the different gene dependencies in different cell lines (Liu, et al., 2019).

### 3.2 TranSynergy architecture

TranSynergy is a transformer boosted deep learning model for the prediction of drug combination synergy. It includes three major components, input dimension reduction component, self-attention transformer component, and output fully connected component (Figure 2). The input features are composed of three vectors. Each vector has 2401 dimensions, forming a 3×2401 matrix. In the matrix, each column corresponds to a gene or protein. The first two vectors are the representations of two drugs. The third vector is the representation of the cell lines. The input dimension reduction component is a single-layer neural network to reduce the dimension of input. The modified transformer component takes the output from the first component and applies a scaled dot product based self-attention mechanism to it. Here, the self-attention is applied to model gene-gene interactions. It is also worth noting that we customize the transformer model by removing the positional encoding layer since the order input feature dimensions should be irrelevant to the final prediction. Then the final output of the predicted synergy score comes from a fully connected neural network whose architecture is the same as that used in DeepSynergy. The hyperparameters used in each component and training process are listed in the supplementary Table 1.

**Figure 2.**
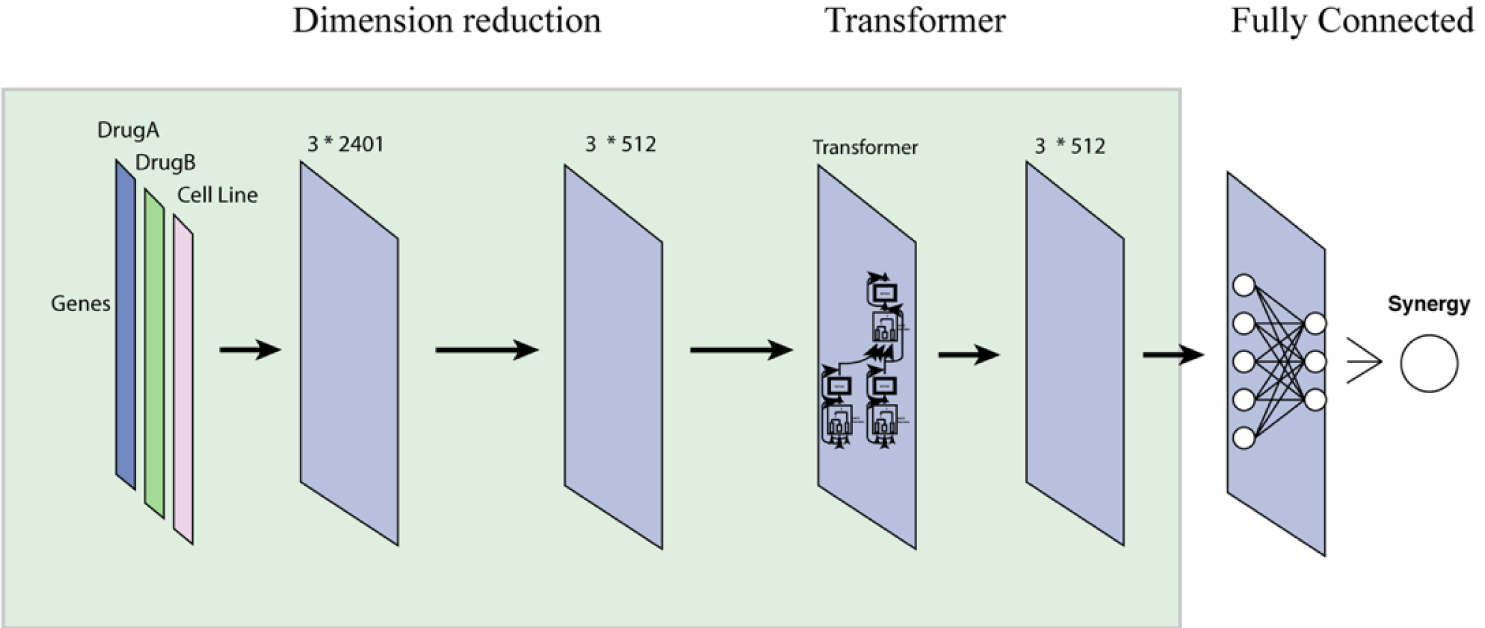
The architecture of TranSynergy. The input features include vector representations of Drug A, Drug B, and cell line vector, respectively. The first input dimension reduction component reduces the input dimension from 3*2401 to 3*512. The second component is a scaled dot product self-attention transformer. The third component is a fully connected neural network with the same architecture as that in DeepSynergy.

### 3.3 TranSynergy outperforms state-of-the-art model and shows superior performance across different cell lines

To test the performance of TranSynergy architecture and compare it with the state-of-the-art models, we used the same dataset in deepSynergy study. 80% of data are used in the training and 20% are held out to test model performance, and all drug combinations in test datasets are unseen in the training dataset. To make an apple-to-apple comparison, we use the same drugs and cell lines vector representation for both two deepSynergy architecture (fully connected only) and TranSynergy architecture. TranSynergy significantly outperforms the deepSynergy model in the synergy score prediction (Table 1). The accuracy improves 5.5% and 8.1% when measured by Pearson’s correlation and Spearman’s correlation, respectively. It is noted that the performance when using NetExpress score is comparable to that using the gene dependency. It is not surprising because NetExpress score is correlated with the gene dependency. Because the gene dependency data are not always available, it is possible for us to simulate the gene dependency using the NetExpress score, and achieve the comparable performance. It is noted that genomics features of cell lines (mutations and CNVs) were not used in both TranSynergy and DeepSynergy. They could provide additional information that is relevant to the drug mode of action, especially for targeted therapy. It will be interesting to incorporate genomics features into TranSynergy in the future.

**Table 1.** Performance comparison of TranSynergy with fully connected neural network (deepSynergy) model.

| Models | Drug features | Cell line features | Pearson's Correlation | Spearman's Correlation |
| --- | --- | --- | --- | --- |
| TranSynergy | Network propagated drug target profile | Gene dependencies | **0.770±0.017** | **0.746±0.011** |
|  |  | NetExpress | 0.764±0.022 | 0.739±0.012 |
| deepSynergy | Physicochemical properties, toxicophores and fingerprints | Gene expression | 0.730±0.020 | 0.690±0.018 |

We further explored the performance of TranSynergy model on 6 different tissues, colon, breast, melanoma, ovarian, prostate and lung. The average Pearson correlation coefficient between ground truth synergy score and predicted scores are 0.767 for colon cancer cells, 0.739 for breast cancer cells, 0.757 for melanoma, 0.797 for ovarian cancer cells, 0.665 for prostate cancer cells, and 0.803 for lung cancer cells, respectively (Figure 3A-B). The performance of TranSynergy across tissues are consistent except the prostate. VCAP is the only prostate cancer cell line, which has the lowest Pearson correlation coefficient for tissues included in our data. VCAP has the second lowest Pearson correlation coefficient and Spearman correlation coefficient across all cell lines. The performance of TranSynergy across cell lines ranges from 0.591 to 0.894 for Pearson correlation coefficient, and from 0.592 to 0.891 for Spearman correlation coefficient (Figure 3C-D), respectively.

**Figure 3.**
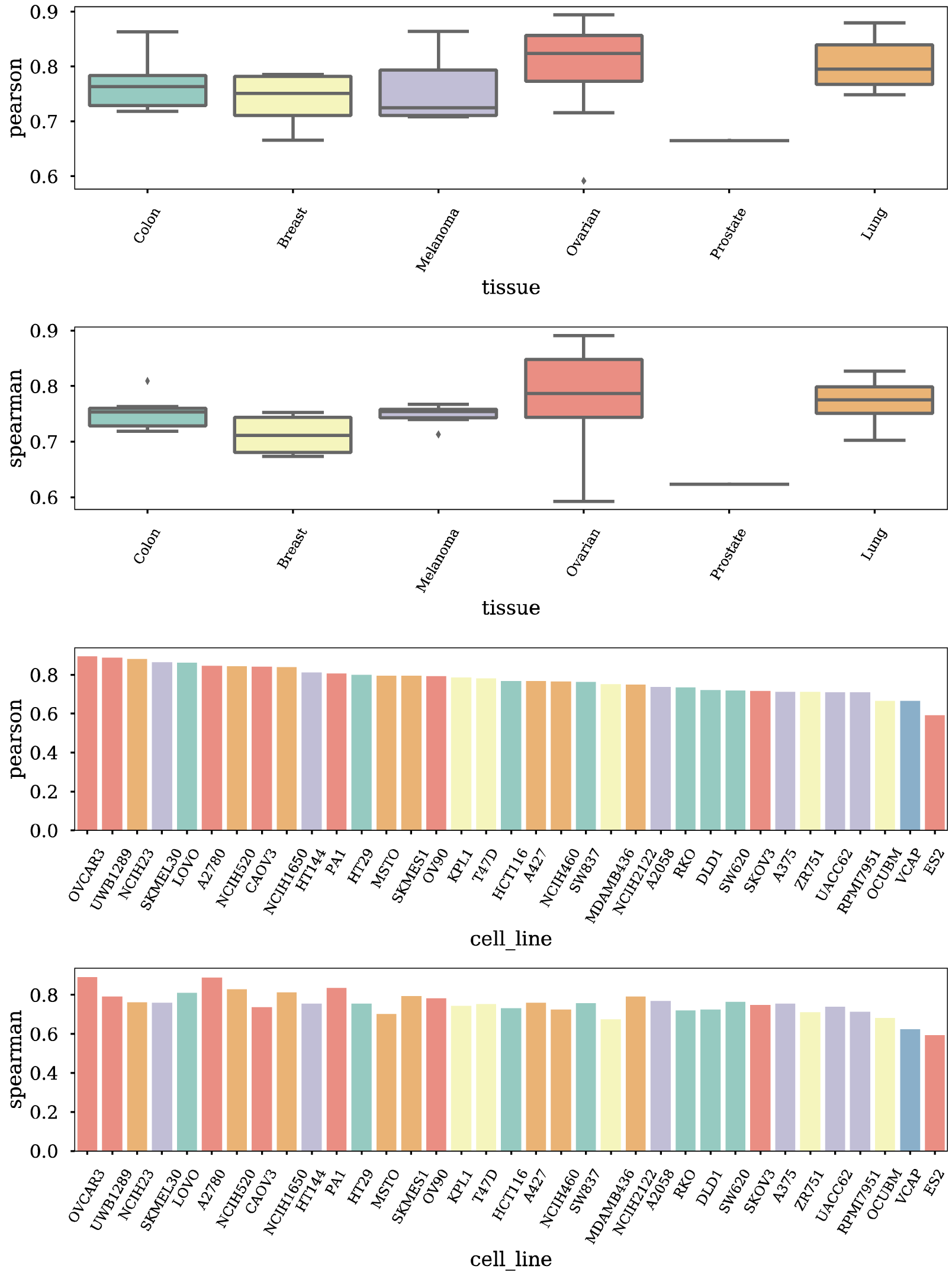
TranSynergy performance in 6 different tissues and 35 cell lines. Box plot of A) Pearson correlation coefficients and B) Spearman correlation coefficients of ground truth synergy scores and TranSynergy model predicted synergy scores in 6 different tissues. Bar plot of C) Pearson correlation coefficients and D) Spearman correlation coefficients of ground truth synergy scores and TranSynergy model predicted synergy scores in 35 cell lines. The color denotes the tissue each cell line belongs to.

### 3.4 Mechanism-driven drug and cell line representations and model architecture are critical for superior model performance

We introduced two strategies for cell line vector representation based on gene dependency and NetExpress score which combines the gene expression and gene-gene interaction to surrogate the gene dependency. To demonstrate the importance of these kinds of mechanism-driven representations, we compared their performance with conventionally used gene expression only representation on both TranSynergy and DeepSynergy architectures. As shown in Table 2, we found that models that are based on the gene dependency and NetExpress score perform better than models with gene expression only representation. Because both gene dependencies and NetExpress scores characterize the dependencies of cells on each gene for different cell lines, they provide more mechanism information on cellular response to drugs than gene expression only.

**Table 2.** Ablation study of TranSynergy model.

| Drug features | Models | Cell line features | Pearson's Correlation | Spearman's Correlation |
| --- | --- | --- | --- | --- |
| Network propagated drug target profile | With self-attention | Gene dependencies | **0.770±0.017** | **0.746±0.011** |
|  |  | NetExpress | 0.764±0.022 | 0.739±0.012 |
|  |  | Gene expression | 0.731±0.028 | 0.713±0.023 |
|  | Without self-attention | Gene dependencies | 0.734±0.022 | 0.722±0.020 |
|  |  | NetExpress | 0.732±0.021 | 0.721±0.019 |
|  |  | Gene expression | 0.723±0.021 | 0.712±0.020 |
| Observed drug target profile | With self-attention | Gene dependencies | 0.635±0.048 | 0.634±0.025 |
|  | Without self-attention |  | 0.632±0.039 | 0.634±0.038 |

To investigate whether the PPI network propagation step is essential for the model performance, we trained the TranSynergy model with only observed drug target information. The observed drug target vector is a binary 2401-dimension vector that indicates whether a drug physically binds to the corresponding protein. The cell line vector representation used in this comparison is that based on the cell line dependency. We also trained a fully connected neural network model with the same data. Models with the network propagated drug target vector representation show superior performance to those with observed drug target information (Table 2).

Finally, the self-attention that simulates gene-gene interactions plays a critical role in the model performance. As shown in Table 2, regardless of drug and cell line features used, the model with the self-attention consistently outperforms those without the use of self-attention.

### 3.5 Shapley Additive Gene Set Enrichment Analysis reveals novel biological pathways associated with synergistic drug combinations

We propose to use Shapley Additive Gene Set Enrichment Analysis (SA-GSEA) to determine biological pathways associated with synergistic drug combines (see methods for details). Shapley additive value is a powerful way to characterize the feature importance (Lundberg and Lee, 2017). Because each of features in TranSynergy corresponds to a gene, the Shapley value essentially indicates the importance of gene for the synergy prediction. As examples, we explored seven samples in the testing dataset, which have high synergistic scores and are accurately predicted (Table 3 and Supplementary Figures 1-16). Both two best TranSynergy models harbor propagated drug target features. One of them uses NetExpress scores as cell line features, while the other represents cell line with gene dependencies profiles.

**Table 3.** The most important pathways revealed by SA-GSEA.

| Drug combination<br>(drug 1, drug 2) | Cell line | Targets with high SHAP values |  | The most enriched gene set in the cell line |  |
| --- | --- | --- | --- | --- | --- |
|  |  | Drug 1 | Drug 2 | GD | NetExpress |
| BEZ-235,<br>ERLOTINIB | T47D | Targets in<br>PI3K/AKT/mT<br>OR pathway | EGFR | Upregulated<br>genes in RAF<br>overexpressed<br>cells | Downregulated<br>genes in STK33<br>knock-down<br>cells |
| BEZ-235,<br>MK-4827 | T47D | Targets in<br>PI3K/AKT/mT<br>OR pathway | PARP1/2 | Upregulated<br>genes in RAF<br>overexpressed<br>cells | N/A |
| BEZ-235,<br>L-778123 | T47D | Targets in<br>PI3K/AKT/mT<br>OR pathway | FPTase/<br>GGPTase | Upregulated<br>genes in RAF<br>overexpressed<br>cells | Downregulated<br>genes in PRC2-SUZ12 knock-down cells |
| BEZ-235,<br>DINACICLIB | T47D | Targets in<br>PI3K/AKT/mT<br>OR pathway | CDKs | Upregulated<br>genes in RAF<br>overexpressed<br>cells | Downregulated<br>genes in PRC1-BMI knock-down cells |
| ETOPOSID,<br>MK-8669 | CAOV3 | TOP2A/TOP2B | mTOR | Upregulated<br>genes in JAK2<br>knock-down<br>cells | N/A |
| PACLITAXEL,<br>DASATINIB | MSTO-211H | Tubulin/microtu<br>bule associated<br>proteins | BCRABL, SRC,<br>Ephrins and<br>GFR | Upregulated<br>genes in ESR1-cells | Upregulated<br>genes in BMI1<br>knock-down<br>cells |
| ABT-888,<br>DASATINIB | MSTO-211H | PARP1/2 | BCRABL, SRC,<br>Ephrins and<br>GFR | Upregulated<br>genes in SNF5<br>know-down<br>cells | Downregulated<br>genes in PRC1-BMI knock-down cells |

The drug pairs in Table 3 show high synergy score in three cell lines, T47D, CAOV3 and MSTO-211H. For the synergistic drug pairs in T47D, BEZ-235 is one of the notable drugs. The BEZ-235 can inhibit PI3K/AKT/mTOR pathway, which is deregulated in breast cancer (Verret, et al., 2019). In the SA-GSEA of cell line gene dependencies profile, RAF oncogenic signature is significantly enriched. It is established that RAS/RAF/MEK/ERK pathway and PI3K/AKT/mTOR pathway are closely interconnected components and form feedback loops in breast cancer (Saini, et al., 2013). The SA-GSEA of cell line NetExpress scores reveals additional pathways associated with the synergistic drug combination, including STK33, PRC2-SUZ12, and PRC1-BMI oncogenic signatures. STK33 can interact with PI3K/AKT/mTOR pathways (Castellano and Downward, 2011; Zhou, et al., 2020). PRC2-SUZ12 pathway is known to play a crucial role in DNA damage-repairing process (Bhoumik, et al., 2005; Campbell, et al., 2013; Williams and Schumacher, 2016). Besides, PI3K/AKT/mTOR pathway activate transcription by reducing trimethylation of promoter-associated Histone H3 Lys27 (H3K27me3), which is regulated by PRC2 (Spangle, et al., 2017). This may be one explanation of the observed cell growth stalling after PI3K/AKT/mTOR inhibition. Similarly, AKT can phosphorylate BMI, which reduces the transcriptional silencing caused by BMI (Spangle, et al., 2017). This may also suggest the underlying mechanism following PI3K/AKT/mTOR pathway inhibition.

The drug combination of ETOPOSIDE and MK-8669 is synergistic in CAOV3 cancer cell line. ETOPOSIDE targets TOP2A and TOP2B, two topoisomerase components. The inhibition on them is believed to cause DNA double strand break. MK-8669 targets mTOR, a crucial component in PI3K/AKT/mTOR pathway. Besides, TUBB also shows a surprising high importance in the SA-GSEA of MK-8669 drug targets. This indicates that inhibition of mTOR or TUBB combining with DNA damage can have a synergistic effect. In the SA-GSEA of cell line features, JAK2 oncogenic signature is enriched in the ranked gene list based on their Shapley values. This might suggest that JAK2 is novel pathway affected by this drug combination therapy (Bartalucci, et al., 2013).

For synergistic drug pair in MSTO-211H cell line, one of the drugs is DASATINIB. The DASATINIB targets BCRABL, SRC, Ephrins and GFR. For the combination of PACLITAXEL and DASATINIB in MSTO-211H cell line, the PACLITAXEL targets tubulin and microtubule associated proteins. From the SA-GSEA of cell line genes, genes in the ESR1 and BMI1 oncogenic signature gene sets were top ranked. SRC activates estrogen receptor *α* proteolysis in some cancers, thus inhibition of SRC can alters cancer cell response to other therapies (Chu, et al., 2007). Investigation has found that BCRABL can enhance BMI1 expression (Bhattacharyya, et al., 2009). BMI1 is also shown to be involved in DNA-damage-repair process (Ginjala, et al., 2011). The inhibition of these pathways combining the restrain on mitosis can have high anti-tumor activities in Mesothelioma cancer. For the combination of DASATINIB and ABT-888 in MSTO-211H, besides BCRABL-BMI interaction mentioned before, SNF5 pathway is also ranked on the top. It worth mentioning that both SNF5 and PARP1/2 play key roles in the DNA repair process (Javle and Curtin, 2011; Ribeiro-Silva, et al., 2019).

### 3.6 Novel drug combination prediction

We predicted the synergy score of novel drug pairs-cell line samples. For 3650 novel samples, Table 4 lists top 10 samples which have higher cell line-wise z-scores (Supplementary Table 2). It is noted that Etoposide, which targets TOP2A and TOP2B, is one of drug components in the drug pairs which are synergistic on CAOV3 cancer cell line. It is consistent with the results of SA-GSEA in which the target combination of TOP2A, TOP2B and TUBB shows high feature importance. Two samples show synergistic effect on OV90, another ovarian cancer cells. For ETOPOSIDE and VINORELBINE combination, drugs also target TOP2A, TOP2B and TUBB. For PACLITAXEL and VINORELBINE, both drugs target tubulin or microtubule associated proteins. Besides, DEXAMETHASONE exists in synergistic drug combinations in OCUBM, SKMES1and KPL1.

**Table 4.** Predicted most synergistic novel drug pairs.

| Drug combination | Cell line | Predicted Synergy score | Z-score |
| --- | --- | --- | --- |
| DEXAMETHASONE and ETOPOSIDE | CAOV3 | 94.520215 | 3.07259459 |
| ETOPOSIDE and METFORMIN | CAOV3 | 70.556332 | 2.21009253 |
| ETOPOSIDE and SN-38 | CAOV3 | 69.380475 | 2.16777138 |
| DEXAMETHASONE and VINBLASTINE | OCUBM | 38.6095235 | 1.94930657 |
| DEXAMETHASONE and PACLITAXEL | SKMES1 | 37.9697725 | 1.91635083 |
| ETOPOSIDE and VINOELBINE | OV90 | 33.844834 | 1.89400207 |
| PACLITAXEL and VINOELBINE | OV90 | 32.5451685 | 1.81387084 |
| ETOPOSIDE and VINBLASTINE | CAOV3 | 56.799048 | 1.71494381 |
| DEXAMETHASONE and VINBLASTINE | KPL1 | 37.8084735 | 1.6825975 |
| ETOPOSIDE and PACLITAXEL | CAOV3 | 53.1252895 | 1.58271882 |

## 4 Discussion

In this study, we presented a novel deep learning model TranSynergy for the synergistic prediction and mechanism deconvolution of drug combinations. We demonstrated that network propagated drug target profile, which indicates both drug-target protein interaction and drug effect on non-targeted proteins, is crucial for the comprehensive representation of drug features. We also showed that gene essentiality in the cancer cells, whether experimentally or computationally determined, is a better cell line representation than raw gene expression profile. Due to the limited data size, we only selected the minimum number of genes for the representation of drugs and cell lines, which includes only drug target genes and annotated cancer-related genes. Too many input features could cause the model to suffer overfitting problems during the training due to the curse of dimensionality (Clarke, et al., 2008; Danaee, et al., 2017). The performance can be improved further when more data are available or by utilizing unlabeled data sets so that more genes can be included into the representations.

Even though multiple high throughput drug combination screening dataset is available, we still face the inconsistency problem in both the experimental and quantification methods for the determination drug combinations effects. Firstly, the combinational spaces for the drug doses used to generate the drug dose matrix vary in different studies. Secondly, several distinguished methods are proposed to calculate the expected drug combination effect from experimental data, such as combination index (CI)-isobologram equation (Chou and Talalay, 1983; Chou, 2010; Chou and Talalay, 1984), Bliss independence (BI) method (Berenbaum, 1978; Bliss, 1939; Greco, et al., 1995), and Loewe Additivity (LA) model (Loewe, 1953; Loewe and Muischnek, 1926). The calculated drug synergy scores are not the same when different quantification methods are utilized. To further improve the data quality, it is necessary to develop new methods to harmonize different data sets.

Deep learning-based computational models have made promising breakthroughs in many biomedical areas. Interpretation of deep learning models becomes critical to overcoming the skepticism of it being a black-box (Ching, et al., 2018). Recently, many methods have been proposed, such as input perturbation methods (Heaton, et al., 2017; Zhou and Troyanskaya, 2015), backpropagation based methods (Springenberg, et al., 2014), and the calculation of SHAP values (Lundberg and Lee, 2017). We carefully design the input features so that each feature dimension is corresponding to a gene. With the SHAP values of the gene-wise input feature, we extract the information on the effect of drug-target interactions and gene-gene interactions on the cancer cell response. This potentially provides a method to study the underlying mechanism of the multi-targeted drug combinations therapy.

Drug combinations can be a more efficient therapeutic strategy for cancer by targeting multiple proteins to defer the rapid emergence of drug resistance. The exploration of effective and synergistic drug combinations is hindered by the costly and time-consuming experimental preclinical investigation. Computational methods can be a cheaper and faster alternative approach to facilitate the development of drug combination therapy for cancer patients (Yin, et al., 2018). Nowadays, more emphasis is put on personalized medicine, which requires the consideration of the heterogeneity of each patient’s cancer types and genomics information to find more efficient therapy. Given a large amount of data for patients’ genome information, development of an accurate and interpretable computational model is critical for the realization of personalized medicine. Mechanism-driven machine learning as demonstrated in this study is a promising direction to address challenges in precision medicine of combination therapy.

