## Supplemental Table 1 for "TranSynergy: Mechanism-Driven Interpretable Deep Neural Network for the Synergistic Prediction and Pathway Deconvolution of Drug Combinations"

Supplementary Table 1. The hyperparameters of TranSynergy model with the best performance.

|  |  | TranSynergy |
| --- | --- | --- |
| Dimension reduction Layer | Number of hidden unit | 512 |
|  | Embedding output dimension | 512*3 |
| Transformer | Embedding dimension | 400 |
|  | Feed-forward hidden unit | 200 |
|  | Number of heads | 1 |
|  | Output dimension | 512*3 |
| Fully connected | Number of hidden unit | [2000, 1000, 1] |
| Dropout | | 0.2 |
| Learning rate | | 0.0001 |
| Weight decay | | 0.00001 |
